# Integrated global analysis in spider-flowers illuminates features underlying the evolution and maintenance of C_4_ photosynthesis

**DOI:** 10.1101/2022.10.06.511187

**Authors:** Wei Zhao, Jun Li, Xingchao Sun, Qiwei Zheng, Wei Hua, Jun Liu

**Affiliations:** Key Laboratory of Biology and Genetic Improvement of Oil Crops, Ministry of Agriculture and Rural Affairs, Oil Crops Research Institute of the Chinese Academy of Agricultural Sciences, Wuhan 430062, China; Hubei Hongshan Laboratory, Wuhan 430070, China

**Keywords:** convergent evolution, reference genome, C_4_ biology, model plant, *Gynandropsis gynandra*

## Abstract

The carbon concentrating mechanism—C_4_ photosynthesis—represents a classic example of convergent evolution. While how this important trait originated and evolved remains largely enigmatic. Here we present a high-quality chromosome-scale annotated genome assembly of the spider-flower *Gynandropsis gynandra*, a valuable leafy vegetable crop and medicinal plant that has also been recognized as an emerging C_4_ model species. Repetitive elements occupy up to 71.91% of its genome, and over half are LTR-RTs derived from recent bursts, contributing to genome size expansion. Strikingly, LTR-RT explosion also played a critical role in C_4_ evolution by altering expression features of photosynthesis-associated genes via preferential insertion in promoters. Synteny analysis in the *Cleome* genus unveils that an independent species-specific whole-genome duplication in *G. gynandra*, which we name Gg-α, occurred after divergence from its close relative C_3_ plant *Tarenaya hassleriana*. Integrated multi-omics profiling demonstrates that Gg-α, gene family expansion, recent LTR-RT amplification and more recent species-specific tandem duplication events have all facilitated the evolution of C_4_ photosynthesis, revealing uniqueness of C_4_ evolution in this lineage. Moreover, high leaf vein density and heat stress resilience are associated with shifted gene expression patterns. Altogether, this mode of C_3_-to-C_4_ transition yields new insights into evolutionary convergence of a complex plant trait.

**One-sentence summary:** We present a high-quality chromosome-scale genome assembly for dicot C_4_ model crop *Gynandropsis gynandra* and find an independent evolutionary trajectory from C_3_ to C_4_ photosynthesis in *Cleome* genus.

## INTRODUCTION

Photosynthesis is the basis of most life forms on the planet. C_4_ photosynthesis represents a remarkable convergent innovation that results from a series of anatomical and biochemical modifications to the ancestral C_3_ photosynthetic pathway, which together function to increase CO_2_ concentration around the enzyme RuBisCO, thereby reducing photorespiration and enhancing photosynthetic efficiency (Schlüter and Weber, 2020). Most, if not all, C_4_ species, which possess a distinctive leaf structure characterized by Kranz anatomy, are typically classified into three metabolic subtypes based on the enzymes used to decarboxylate C_4_ acids in their bundle sheath cells (BSC): NADP-ME, NAD-ME and PEP-CK (Zhao et al., 2022; Furbank, 2011). C_4_ plants usually have higher photosynthetic capacity and higher nitrogen and water-use efficiencies than their C_3_ relatives and thus tend to be more productive (Sage, 2004; Sage et al., 2012). In particular, C_4_ photosynthesis outperforms the ancestral C_3_ state under sunny, hot and dry circumstances, which are projected to become more prevalent with global climate changes (Hibberd et al., 2008; Wang et al., 2021).

The introduction of dual-celled C_4_ photosynthesis into C_3_ plants is proposed to be a promising strategy to sustainably meet the rising food, fuel and feed demands worldwide (Wang et al., 2016; Ermakova et al., 2020). This goal requires a profound understanding of the origin, genetic architecture and developmental features of C_4_ syndrome as compared to C_3_. Several C_4_ monocots, including maize (*Zea mays*), sorghum (*Sorghum bicolor*) and millets (*Setaria viridis* and *Setaria italica*), have been used as model plants for this purpose (Paterson et al., 2009; Mamidi et al., 2020; Bennetzen et al., 2012; Jiao et al., 2017). However, these models share various disadvantages including relatively large size, complex genomes, long life cycles and low transformability (Brown et al., 2005). Fortunately, a mini foxtail millet mutant, *xiaomi*, has recently been established as a new NADP-ME subtype C_4_ model system (Yang et al., 2020). However, all current C_4_ model plants are monocotyledonous species, and there are differences in C_4_ attributes of dicots and monocots, including vein patterning, Kranz anatomy morphogenesis, regulation of C_4_ photosynthetic enzymes and development of C_4_ capacity (Ermakova et al., 2020; Schlüter and Weber, 2020). Therefore, it is vital to develop a dicot C_4_ model plant and would furthermore be advantageous to develop a model system in NAD-ME or PEP-CK subtypes of C_4_ plants. Investigation of these C_4_ model organisms would accelerate systematic understanding of C_4_ biology and facilitate synthetic engineering of the C_4_ pathway into contemporary C_3_ crops.

The dicot *Gynandropsis gynandra* (common names: spider-flower, African cabbage, bastard-mustard and cat’s-whiskers) is an important traditional C_4_ crop native to Asia and Africa, currently cultivated around the world (Oshingi et al., 2019). The leaves and seeds of *G. gynandra* are rich in proteins, vitamins, minerals and other beneficial health compounds with antioxidant, anti-inflammatory and anti-microbial properties, and *G. gynandra* has thus been widely used as a leafy vegetable or medicinal plant (Moyo and Aremu, 2021). As a C_4_ species, *G. gynandra* displays high photosynthetic efficiency and adaptation to severe environmental stresses such as high temperature, water deficit or high salinity (Sogbohossou et al., 2018). Importantly, it is a diploid crop characterized by relatively short life cycle, small size, simple growth requirements, prolific seed production, self-compatibility and autogamy. Besides, *G. gynandra* has a rich diverse germplasm that provides valuable genetic material for dissecting traits of interest and breeding (Sogbohossou et al., 2019). These advantages, coupled with its efficient genetic transformation system, make this plant an ideal model plant for C_4_ biology (Newell et al., 2010).

*G. gynandra* belongs to family of Cleomaceae, the sister clade to Brassicaceae, and is phylogenetically closest to the dicot C_3_ model plant *Arabidopsis thaliana*, enabling the utilization of molecular resources and tools developed for *Arabidopsis* in *G. gynandra* and facilitating knowledge transfer (Marshall et al., 2007; Koteyeva et al., 2011; Patchell et al., 2014; Hoang et al., 2022). Moreover, *G. gynandra*, together with the congeneric C_3_ species *Tarenaya hassleriana* (common names: spider-flower, pinkqueen and grandfather’s-whiskers), provides an invaluable genetic platform for comparative studies of many intriguing biological phenomena including the evolutionary trajectory and developmental progression from C_3_ to C_4_ (Bräutigam et al., 2011; Külahoglu et al., 2014; Bayat et al., 2018; Hüdig et al., 2022). However, there is no published reference genome for *G. gynandra*, which hinders its use for fundamental biological discovery, technological advances and applications for genetic engineering. To address this gap, we generated a high-quality chromosome-level reference genome assembly of *G. gynandra*, establishing it as a C_4_ model system for dicotyledonous and NAD-ME subtype plants. Our comprehensive comparative genomic and transcriptomic analyses between *G. gynandra* and *T. hassleriana* using new and existing data not only help explain why C_4_ photosynthesis failed to evolve in *T. hassleriana* but also demonstrate the flexibility of the convergent evolution of this ecologically important but complex trait. The availability of this reference-grade genomic resource, together with its amenability to genetic transformation, makes *G. gynandra* an ideal model system for understanding the unique C_4_ biology and facilitating efforts toward C_4_-aimed crop engineering.

## RESULTS

### Chromosome-scale assembly and annotation of *G. gynandra* genome

Based on *k*-mer analysis, the dicotyledonous C_4_ plant *G. gynandra* (2n=2x=34) had an estimated genome size of approximately 997.61 Mb with low heterozygosity (0.13%) but high repetitive sequence content (79.72%) (Supplemental Figure S1 and Supplemental Data Set S1). To construct a reference-grade genome for *G. gynandra*, we employed an optimized strategy combining long-read Oxford Nanopore Technology (ONT), short-read Illumina sequencing and high-throughput chromatin conformation capture (Hi-C) for chromosome scaffolding (Supplemental Figure S2). A total of 144.37 Gb (~180 ×) of ONT long sequences with an N50 read length of 24.57 kb were generated (Supplemental Figure S3 and Supplemental Data Set S2). The ONT long reads were *de novo* assembled into contigs, followed by polishing with both ONT and Illumina reads. To further anchor and orient the contigs onto chromosomes, we prepared Hi-C libraries from young leaves to construct chromatin interaction maps, generating 130 Gb (~162 ×) paired-end reads (Supplemental Data Set S3). This resulted in a final assembly of 984.21 Mb comprising 109 scaffolds, which represented 98.66% of the estimated nuclear genome (Table 1 and Supplemental Data Set S4). The contig N50 and scaffold N50 were 11.43 Mb and 51.02 Mb, respectively. Sequences of 909.61 Mb covering 171 contigs were assigned to seventeen pseudo-chromosomes (Chr1– Chr17; Figure 1A and Table 1), which accounted for 92.42% of the assembly. The pseudo-chromosomes of *G. gynandra* ranged from 41.13 to 72.98 Mb in size (Supplemental Data Set S5). Of note, a 55.56 Mb contig, which covered 98.7% of the intact Chr5, constituted a near complete, telomere-to-telomere assembly of one *G. gynandra* chromosome.

**Table 1.**
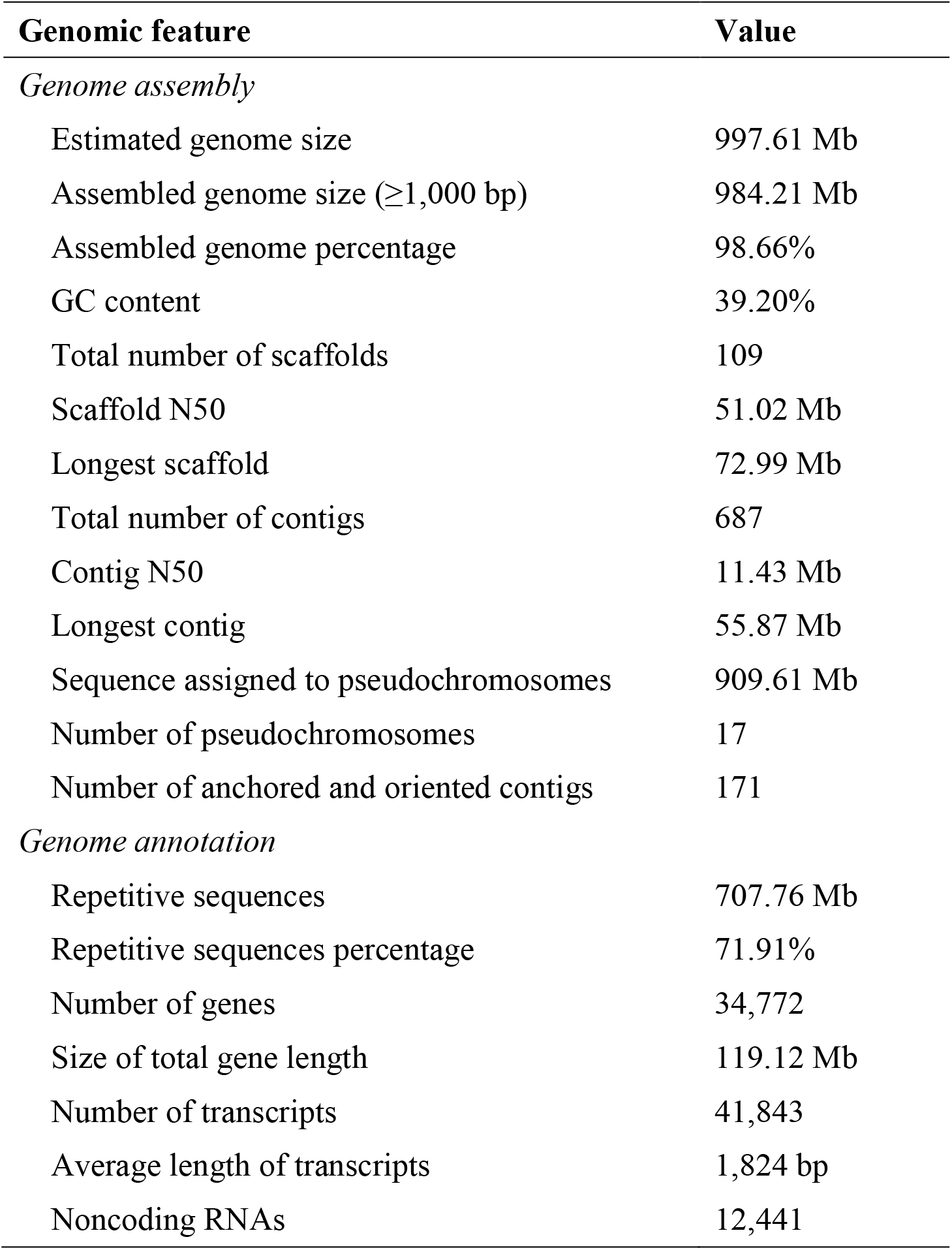
Global statistics of *G. gynandra* genome assembly and annotation.

**Figure 1.**
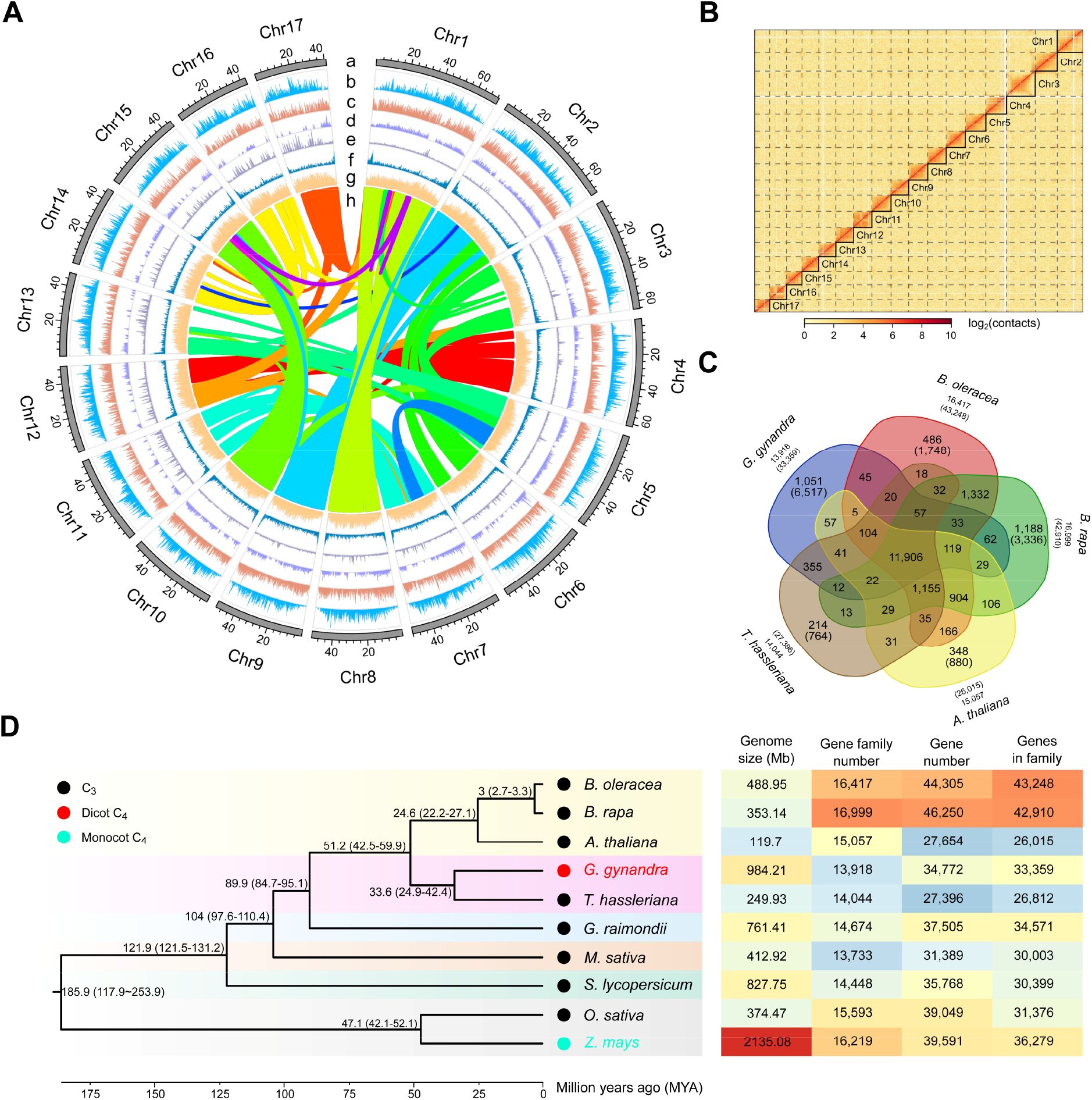
The high-quality assembly, genome features and evolutionary analysis of *G. gynandra*. **(A)** An overview of the genomic characteristics of *G. gynandra*. The outermost annotations number successively the seventeen assembled chromosomes in descending order of size. Tracks from outer to inner circles depict length (Mb) of each chromosome (a), *Gypsy* retrotransposon distribution (b), *Copia* retrotransposon distribution (c), DNA transposon distribution (d), tandem repeats (e), gene density (f), GC content (g) and curve lines in the interior link the syntenic regions that have been retained presumably since the last whole-genome duplication event (h). Sliding window size is 200 kb. **(B)** Genome-wide Hi-C heatmap of the *G. gynandra* genome. Heatmap shows the diagonal pattern for strong Hi-C interactions within intra-chromosome of *G. gynandra*. The X-axis and Y-axis indicate the order positions of scaffolds on corresponding pseudo-chromosomes. The color bar denotes the interaction frequencies of the Hi-C links. **(C)** Venn diagram illustrating the shared orthologous groups (orthogroups) among five species: *G. gynandra, T. hassleriana, B. rapa, B. oleracea* and *A. thaliana*. Each number represents the number of gene families shared among genomes. The number listed in parentheses is the total gene number among the orthogroups. **(D)** Phylogenetic relationships of the two Cleomaceae and eight other species used in the study (*B. oleracea, B. rapa, A. thaliana, G. raimondii, M. sativa, S. lycopersicum, O. sativa* and *Z. mays*). The phylogenetic tree was constructed from single-copy orthologs of these species. Lineage divergence time is indicated at each branch point. The photosynthesis subtype of each species is marked by colored dots at each node. The genome size, gene family number, gene number and number of genes belonging to gene families for each species are listed in the heatmap at right.

The assembly quality of the *G. gynandra* genome was evaluated by multiple approaches. First, Hi-C interaction matrices for the constructed pseudo-chromosomes visualized as a Hi-C heatmap showed a clear anti-diagonal pattern for intra-chromosomal interactions (Figure 1B). Second, the completeness of assembly as analyzed with Benchmarking Universal Single Copy Orthologues (BUSCO) was 95.6% (Supplemental Data Set S6). Third, RNA-seq reads obtained from six representative tissues of flower bud, flower, leaf, root, stem and silique were mapped back onto *G. gynandra* genome, to which approximately 96.16–98.33% of the reads could be aligned (Supplemental Data Set S7). Finally, long terminal repeat (LTR) assembly index (LAI), a metric using intact LTRs to evaluate assembly continuity, has been particularly utilized to assess assembly quality of plant genomes with high repetitive sequence content (Ou et al., 2018). The LAI score of *G. gynandra* assembly was 16.46, higher even than that of the *A. thaliana* reference genome (LAI = 15.62). Collectively, these data demonstrate that *G. gynandra* genome assembly is of high quality in contiguity, completeness and accuracy.

By integrating *ab initio*-based prediction, protein homology-based prediction and RNA-seq/Iso-Seq data, we annotated a total of 34,772 protein-coding genes spanning 119.12 Mb regions, representing 12.1% of the *G. gynandra* genome (Table 1). The gene density distribution along each chromosome was uneven, with higher gene density towards the ends of chromosome arms (Figure 1A). The total number of identified transcripts (including splicing variants) was 41,843. Functional analysis predicted 33,100 (95.19%) genes with known functional annotations in public databases, suggesting highly reliable gene prediction (Supplemental Data Set S8). Moreover, we identified 2,359 non-coding RNAs including 120 miRNAs, 1,153 tRNAs and 1,086 rRNAs (Supplemental Data Set S9), and also 2,178 transcription factors (Supplemental Data Set S10). Of all the protein-coding genes, 95.9% was assigned to 13,918 gene families in *G. gynandra*, which was comparable to *T. hassleriana* (97.8% to 14,044 gene families). However, the former had an average of 2.39 genes per family, with the latter possessing 1.91 (Supplemental Data Set S11). Compared with *A. thaliana, B. rapa, B. oleracea* and *T. hassleriana*, 1,051 gene families containing 6,517 genes were specific to *G. gynandra* (Figure 1C). These unique genes were primarily enriched in metabolic pathways such as biosynthesis of amino acids, carbohydrates, lipids, cofactors and vitamins, terpenoids and polyketides, which is consistent with the nutraceutical food and ethnopharmacological medicinal uses for *G. gynandra* (Supplemental Figure S4).

Phylogenetic analysis was performed based on 661 single-copy orthologous genes among ten angiosperm species including eight dicots (*A. thaliana, Brassica oleracea, Brassica rapa, G. gynandra, T. hassleriana, Gossypium raimondii, Medicago Sativa* and *Solanum lycopersicum*) and two monocots (*Oryza sativa* and *Z. mays*). The results revealed that *G. gynandra* was the most closely related to *T. hassleriana*, both of which were adjacent to *A. thaliana* (Figure 1D). Using a reference divergence times of *B. oleracea*–*B. rapa, Z. mays*–*O. sativa* and *A. thaliana*–*O. sativa* obtained from the TimeTree database, the divergence between Cleomaceae and Brassicaceae was estimated as 51.2 million years ago (MYA), with C_4_ *G. gynandra* and C_3_ *T. hassleriana* sharing a common ancestor approximately 33.6 MYA. Remarkably, the genome size of *G. gynandra* (984.21 Mb) is nearly fourfold as large as *T. hassleriana* (249.93 Mb).

### Recent bursts of transposons massively bloated the *G. gynandra* genome, and LTR-RTs facilitated the evolution of C_4_ photosynthesis

By combining *de novo*- and homology-based repeat family identification approaches, we annotated a total of 707.8 Mb repetitive sequences, representing 71.91% of the *G. gynandra* genome (Table 1). 613.5 Mb of sequences (62.33% of the total assembly) were classified as transposable elements (TEs), including 485.73 Mb (79.17% of TEs) of LTR retrotransposons (LTR-RTs) and 118.88 Mb of DNA transposons, whereas their size and proportion were dramatically lower in *T. hassleriana* (Figure 2A and Supplemental Data Set S12). The spatial distribution of LTR-RTs along chromosomes of *G. gynandra* was uneven (Supplemental Figure S5). Its genome was found containing a considerable number of intact LTR-RTs with sequence length up to 10,000 bp, with a peak at around 7,500 bp, while the peak in *T. hassleriana* occurred at approximately 5,000 bp (Figure 2B). Most (79.72%) intact LTR-RT insertion events in *G. gynandra* genome occurred within 1 MYA, with the peak of amplification occurring around 0.16 MYA, in contrast to approximately 0.31 MYA for *T. hassleriana* (Figure 2C). At the superfamily level, very recent amplifications of *Gypsy* retrotransposons occurred approximately 0.14 MYA in *G. gynandra* (Figure 2D). *Gypsy* retrotransposons had the highest distribution density in the centromeric area and the lowest toward telomeric regions, with *Copia* showing a continuous distribution pattern (Figure 1A).

**Figure 2.**
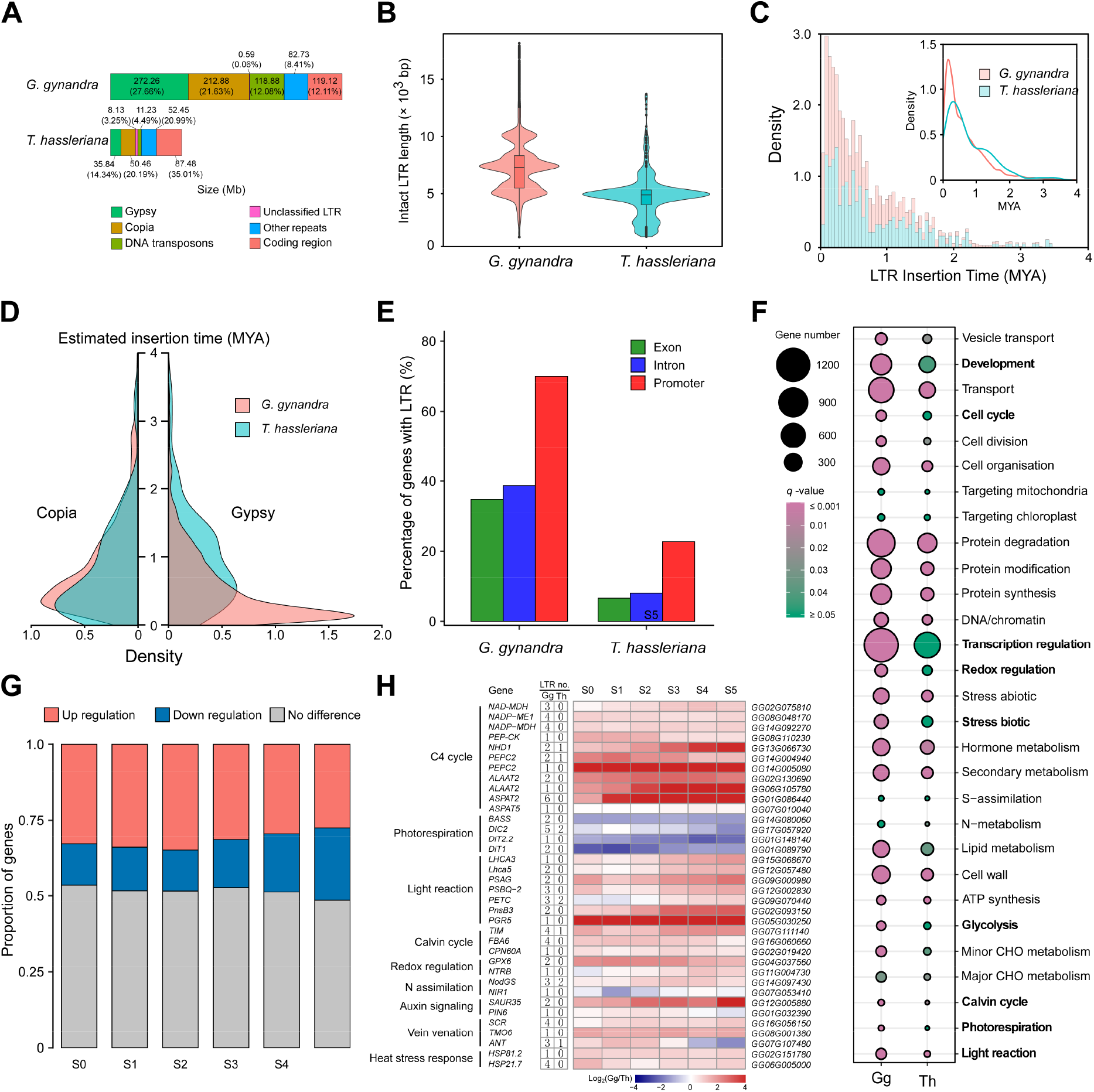
Comparative analysis of TEs in *G. gynandra* and *T. hassleriana* genomes, and the role of LTR-RTs in C_4_ photosynthesis evolution. **(A)** Genomic makeup by category of *G. gynandra* and *T. hassleriana*. The colored blocks indicate the sizes (Mb) of different components, including LTR-RTs (*Gypsy* and *Copia*), DNA transposons, unclassified LTR-RTs, other repeats and coding regions. The number in parentheses is the percentage each taking of the genome. Each number denotes the size (Mb) of each composition. *Gypsy, Copia* and DNA transposons were much more abundant in the *G. gynandra* genome than in *T. hassleriana*. **(B)** Length distribution of intact LTR-RTs in *G. gynandra* and *T. hassleriana*. Boxes within violin plots indicate the first quartile, the median and the third quartile with whiskers extending up to 1.5× inter quartile range (IQR). Outliers are shown as dots, defined as data points outside 1.5× IQR. The *G. gynandra* genome contained much more intact LTR-RTs of over 10 kb in length than *T. hassleriana*. **(C)** Estimated insertion times of LTR-RTs into *G. gynandra* and *T. hassleriana* genomes. The X and Y axes indicate the insertion times and the density of intact LTR-RTs at each time, respectively. *G. gynandra* underwent more extensive LTR-RT explosion than *T. hassleriana* during the last 2 MYA. **(D)** Temporal patterns of *Gypsy* and *Copia* bursts in *G. gynandra* as compared to *T. hassleriana*. The X-axis and Y-axis indicate the density of *Gypsy*/*Copia* and the insertion times, respectively. Despite *G. gynandra* and *T. hassleriana* shared similar density distributions of *Copia*, the former experienced a surge of *Gypsy* insertion very recently. **(E)** Percentage of LTR-RT exon-, intron- or promoter-inserted genes. The 2 kb region upstream of the transcription start site is defined as the promoter region of gene. Genes with LTR-RT insertion in promoter regions are much more abundant than in exons or introns, especially in *G. gynandra*. **(F)** MapMan-Bin enrichment of the LTR-RT promoter-inserted genes in *G. gynandra* and *T. hassleriana*. Annotations of protein sequences with MapMan terms were performed with the online Mercator (https://www.plabipd.de/portal/mercator4). The MapMan4 program was used to conduct the analysis, and the results were visualized with R software. The MapMan terms with clear differences between the two species are highlighted in bold. **(G)** Histograms showing the proportion of selected LTR-RT promoter-inserted genes with up- or down-regulation in *G. gynandra* relative to *T. hassleriana*. The 6,283 genes, which have more LTR-RT insertions in their promoter regions in *G. gynandra* than in *T. hassleriana*, were used for expression analysis. This was performed with the previously deposited RNA-seq data of leaves at six developmental stages (from young to mature, S0 to S5) for these two *Cleome* species (Külahoglu et al., 2014). The gene expression levels were normalized with the upper quartile normalization procedure using the youngest S0 leaf stage of *G. gynandra* as the reference. No difference is defined if the levels of gene expression show lower than a 1.5 fold change between *G. gynandra* and *T. hassleriana*. The X-axis and Y-axis indicate leaf developmental stages and the proportion of genes in each regulation category, respectively. The proportions of up-regulated genes are overall higher than those down-regulated at all the analyzed leaf stages. **(H)** Expression patterns of C_4_ photosynthesis-related pathway genes that have more LTR-RT insertions in their promoter regions in *G. gynandra* than in *T. hassleriana*. Pathways and member genes are indicated at left. The two columns of numbers show how many LTR-RTs were found in the promoter region of each gene in *G. gynandra* and *T. hassleriana*, respectively. The right panel shows the ID of each gene copy in *G. gynandra*. The heatmap illustrates differential expression ratios of the genes between the two species at various leaf developmental stages (S0 to S5). Color reflects fold differences (log_2_ ratios) in gene expression. LTR no., number of LTR-RTs insertion.

Recent massive expansion of LTR-RTs led us to explore their biological relevance. There were 20,094 homologous gene pairs in total between the genomes of *G. gynandra* and *T. hassleriana*. Relative to homologues without LTR-RTs, their presence caused more to be up-regulated (Supplemental Figure S6A). LTR-RTs were found to reside preferentially within promoter regions of protein-coding genes, especially in *G. gynandra* (Supplemental Figure S6B), potentiating their differential roles in regulating gene expressions between the two *Cleome* species. More than 70% of *G. gynandra* genes had LTR-RT insertions in their promoter regions, which were only 23% for *T. hassleriana* (Figure 2E). Functional enrichment analysis of these preferentially inserted genes indicated that pathways of development, cell cycle, transcription regulation, redox regulation, stress-related, glycolysis, Calvin cycle, photorespiration and light reaction were over-represented in *G. gynandra* compared with *T. hassleriana* (Figure 2F). Furthermore, we identified 6,283 genes in *G. gynandra* whose promoter regions contained substantially more LTR-RT insertions than their orthologs in *T. hassleriana* (Supplemental Data Set S13). To assess the consequence of LTR-RT amplification in these genes, their expression levels were analyzed in *G. gynandra* relative to *T. hassleriana*. Moreover, we found that the proportions of genes with up-regulated expression were much greater than those of down-regulated across the six developmental stages, being more pronounced at the early stages (Figure 2G). Notably, the significantly up-regulated genes along the development gradients included a subset of genes associated with C_4_ metabolism (Figure 2H), such as C_4_ cycle (*NAD-MDH, NADP-ME1, NADAP-MDH, PEP-CK, NHD1, ALAAT2* and *ASPAT*), light-harvesting complex (*LhcA3*), photosystem (*PasG* and *PsbQ-2*), cyclic electron flow (CEF, *PnsB3* and *PGR5*), Calvin cycle (*FAB6* and *TIM*), redox regulation (*GPX6* and *NTRB*), N assimilation (*NodGS* and *NIR1*), auxin signaling (*SAUR35* and *PIN6*), vein venation (*SCR, TMO6* and *ANT*), as well as heat stress response (*HSP81*.*2* and *HSP21*.*7*). In contrast, photorespiration genes with more LTR-RTs (*BASS, DIC2* and *DiT*) were severely down-regulated in *G. gynandra*. Additionally, LTR-RT insertions caused few C_4_ metabolism-related genes to be down-regulated as well, such as the copies of *NAD-MDH* (*GG09G088650* and *GG12G084140*), *NADP-ME* (*GG11G010180*) and *PEPC* (*GG02G076930*) (Supplemental Figure S6C). Intriguingly, several C_4_ cycle genes whose promoter regions had less LTR-RTs in *G. gynandra* than in *T. hassleriana* showed much higher levels of expression, such as *NAD-ME* homologues (*GG09G104820* and *GG02G000360*) and *PPDK*. Together, these data reveal that recent large-scale TE bursts are the driving force behind genome size expansion and that LTR-RTs also play a role in the C_4_ evolution of *G. gynandra*.

### Evidence for species-specific whole-genome duplication and tandem gene duplication events in *G. gynandra*

A paleopolyploidization event has been reported for the *Cleome* genus, and *T. hassleriana* underwent a whole-genome triplication (WGT) (Cheng et al., 2013; van den Bergh et al., 2014). However, it remains to clarify whether this event is specific to *T. hassleriana* or shared with *G. gynandra*. We thus analyzed syntenic blocks within the *G. gynandra* genome through intra-genome comparisons, identifying 771 syntenic blocks with 8,801 paralogous gene pairs (Supplemental Data Set S14). We detected multiple duplications in *G. gynandra* based on synteny, with many pairs of paralogous genes. Specifically, chromosomes Chr1, Chr2, Ch3, Chr4, Chr7, Chr10 and Chr14 corresponded closely with Chr8, Chr6, Chr5, Chr12, Chr11, Chr15 and Chr16, respectively (Figures 1A and 3A). Additionally, intra-chromosomal rearrangements, especially inversions, were pervasive in *G. gynandra* such as those near the arm ends of Chr10 and Chr15. Collinearity analysis also showed that chromosome Chr9 and Chr6 shared syntenic regions with Chr2. Likewise, chromosome Chr17 and Chr8 shared syntenic regions with Chr1, and Chr5 was collinear with both Chr3 and Chr13. Overall, these chromosomes displayed high levels of structural variations, suggesting existence of considerable chromosome rearrangements in *G. gynandra*.

For inter-species comparison, we examined genomic synteny between *G. gynandra* and *A. thaliana* and *V. vinifera* chromosomes, resulting in a 2-to-2 syntenic relationship with *A. thaliana*, and 4-to-1 with *V. vinifera* (Figure 3B and Supplemental Figure S7, A and B). Of note, synteny analysis between *G. gynandra* and *T. hassleriana* showed a 2-to-3 syntenic relationship (Supplemental Figure S7C). Given that *V. vinifera* has not undergone genome duplication (Jaillon et al., 2007), and that *A. thaliana* experienced a recent whole-genome duplication (WGD, termed At-α) (Jiao et al., 2011), we thus named the species-specific WGD event in *G. gynandra* and WGT event in *T. hassleriana* Gg-α and Th-α, respectively. To further infer the time of this Gg-α, we calculated the density distribution of *K*_*s*_ (synonymous substitution rate) and D4DTv (distance of fourfold degenerate transversion) values of collinear gene pairs within *G. gynandra, T. hassleriana, A. thaliana* and *B. rapa*. The distribution of *K*_*s*_ values showed that *G. gynandra* had one main peak at *K*_*s*_ of ~0.475 (~22 MYA), which was slightly earlier than *B. rapa* at ~0.313 (~14.7 MYA) and *T. hassleriana* at ~0.390 (~18.4 MYA), and later than *A. thaliana* at ~0.737 (~35 MYA; Figure 3, C and D). The distribution of D4DTv in these four species corroborated time relationships of the WGD or WGT event (Supplemental Figure S7D). About 85.5% of genes in *G. gynandra* genome were duplicated and retained from WGD, which was much higher than the proportion of 63.5% for *T. hassleriana* (Figure 3E). Functional enrichment analysis of these WGD/WGT-derived genes identified pathways such as photosynthesis, ATP synthesis and stress responses as markedly enriched in *G. gynandra* relative to *T. hassleriana* (Supplemental Figure S8A). In addition to WGD, we also identified 610 tandem duplicated genes (TDGs), which were involved in various pathways (Figure 3F). Of importance, the carbon fixation pathway in photosynthetic organisms was exceptionally enriched among *G. gynandra* TDGs, with five duplicated genes including *PEPC2, NAD-MDH*/*mMDH1, GAPC1* and *PDE345*. Their *K*_*s*_ values were all less than 0.1, indicating that these TDGs derived from the very recent duplications after Gg-α (Figure 3G). Except for *PDE345*, the other TDGs exhibited dynamic expression levels from young to mature leaves of *G. gynandra* (Supplemental Figure S8, B and C). Collectively, these results revealed that unlike C_3_ *T. hassleriana, G. gynandra* experienced species-specific WGD (Gg-α) and tandem duplication events, both of which likely facilitated C_4_ photosynthesis evolution.

**Figure 3.**
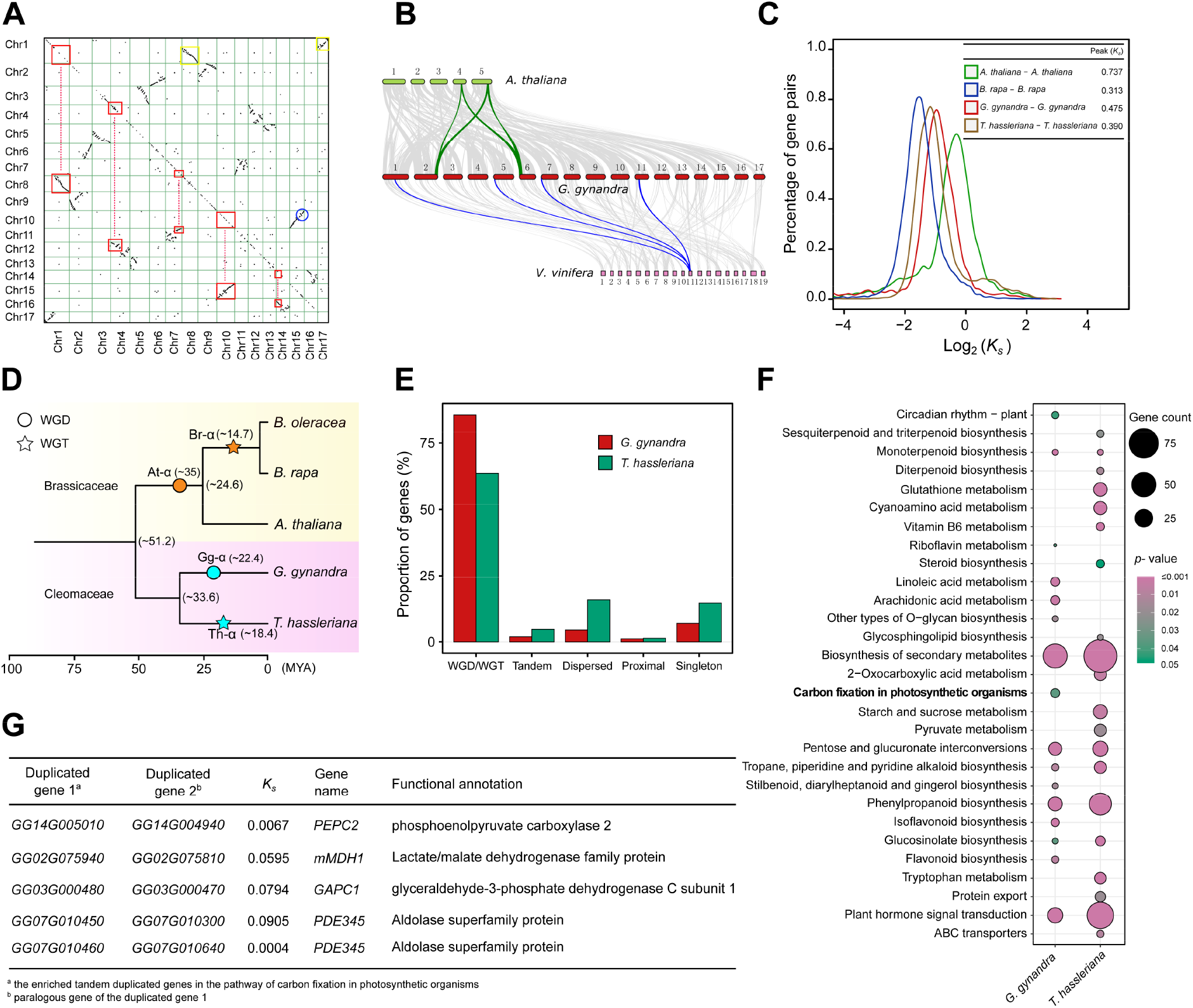
Analysis of WGD and tandem duplications in the *G. gynandra* genome. **(A)** Dotplot figure showing syntenic duplicates within *G. gynandra* genome. The red squares linked by dash lines denote the extensive collinear relationships between chromosomes of *G. gynandra* due to the WGD event. The blue circle represents an intrachromosomal segment inversion. The big and small yellow boxes on Chr1 depict synteny with the chromosomes of Chr8 and Chr17, respectively, suggesting occurrence of potential chromosome breakage and fusion events following WGD. **(B)** Micro-collinearity of *G. gynandra* genome with those of *A. thaliana* and *Vitis vinifera*. The parallel horizontal lines represent the chromosomes of *A. thaliana, G. gynandra* and *V. vinifera* genomes, with the connected grey ribbons indicating syntenic blocks. The 4-to-1 collinear relationship between *G. gynandra* and *V. vinifera* is highlighted by one syntenic set in blue, with one segment in *V. vinifera* traced to four regions in *G. gynandra*. The 2-to-2 collinear relationship between *G. gynandra* and *A. thaliana* is marked by two syntenic sets in green. **(C)** *K*_*s*_ distribution of syntenic orthologs from *G. gynandra, T. hassleriana, A. thaliana* and *B. rapa*. The X and Y axis denotes log_2_(*K*_*s*_) and the percentage of gene pairs in the syntenic blocks, respectively. The values of *K*_*s*_ peaks for each species are showed on the top right corner. **(D)** The evolutionary relationship and paleo-polyploidization events in the five sequenced species of Brassicaceae and Cleomaceae. The paleo-polyploidization and time estimation are indicated on branches of the phylogenetic tree. The yellow and pink background colors represents the Brassicaceae and Cleomaceae families, respectively. The cyan circle and star highlight WGD (Gg-α) and WGT (Th-α) events in *G. gynandra* and *T. hassleriana*, respectively. The divergence time is listed in the brackets. The time scale (MYA) is shown at bottom. **(E)** Gene duplication patterns in the genomes of *G. gynandra* and *T. hassleriana*. The graph indicates the proportion of genes classified by duplication mode relative to total genes of each species. The gene duplication modes were determined by MCScanX, including singletons, dispersed, proximal, tandem and WGD/WGT. WGD and WGT are the major modes of gene duplication for *G. gynandra* and *T. hassleriana*, respectively. **(F)** KEGG enrichments of the tandem duplicated genes in *G. gynandra* and *T. hassleriana*. The size of the circle indicates enriched gene numbers in each pathway, with the color of the circle indicating enrichment *p*-value. Pathways with *p*-value < 0.05 are shown. The “carbon fixation in photosynthetic organisms” pathway is specifically enriched in *G. gynandra* (marked in bold). **(G)** Tandem duplicated genes of the carbon fixation pathway enriched in *G. gynandra*. Duplicated gene paralogs are listed in columns 1–2. *K*_*s*_ values were calculated using the KaKs_Calculator for duplicated gene pairs (Wang et al., 2010). The *K*_*s*_ values of these genes were much lower than the *K*_*s*_ peak of Gg-α (0.475).

### Expanded gene families have contributed to C_4_ trait formation

Closer inspection of gain and loss within gene families showed that 978 and 1,449 ones have undergone expansion and contraction in *G. gynandra*, respectively (Figure 4A). Pfam annotation showed that the genes of expanded families contained protein kinase domain, RING finger, DnaJ domain, PPR repeat, NAD dependent epimerase/dehydratase family, NAD(P)H-binding and AP2 domain (Supplemental Data Set S15). Notably, we identified 221 expanded orthogroups, which were shared by the C_4_ dicot *G. gynandra* and C_4_ monocot *Z. mays* (Figure 4B). These commonly enriched families could be mainly classified into six functional categories: hormone response/signal transduction, gene transcription/protein homeostasis, cell development, leaf development, photosynthetic performance and stress resilience (Figure 4C and Supplemental Figure S9). Specifically, the hormone response/signal transduction category contained a number of genes involved in metabolic processes of auxin, abscisic acid, ethylene or cytokinin, and light signal conduction. Many of these genes also participated in gene transcription/translation or protein homeostasis. Particularly, genes associated with cell and leaf developments were expanded in both C_4_ species, such as cell cycle, programmed cell death, vascular bundles development, chloroplast biogenesis and photomorphogenesis. Expansion of genes in light and in dark reactions could enhance photosynthetic efficiency of C_4_ plants, and gene families involved in resistance to abiotic or biotic stress were also expanded (Figure 4C). These results suggest that the fitness advantages of C_4_ species may have relevance to expansions of defined gene families.

**Figure 4.**
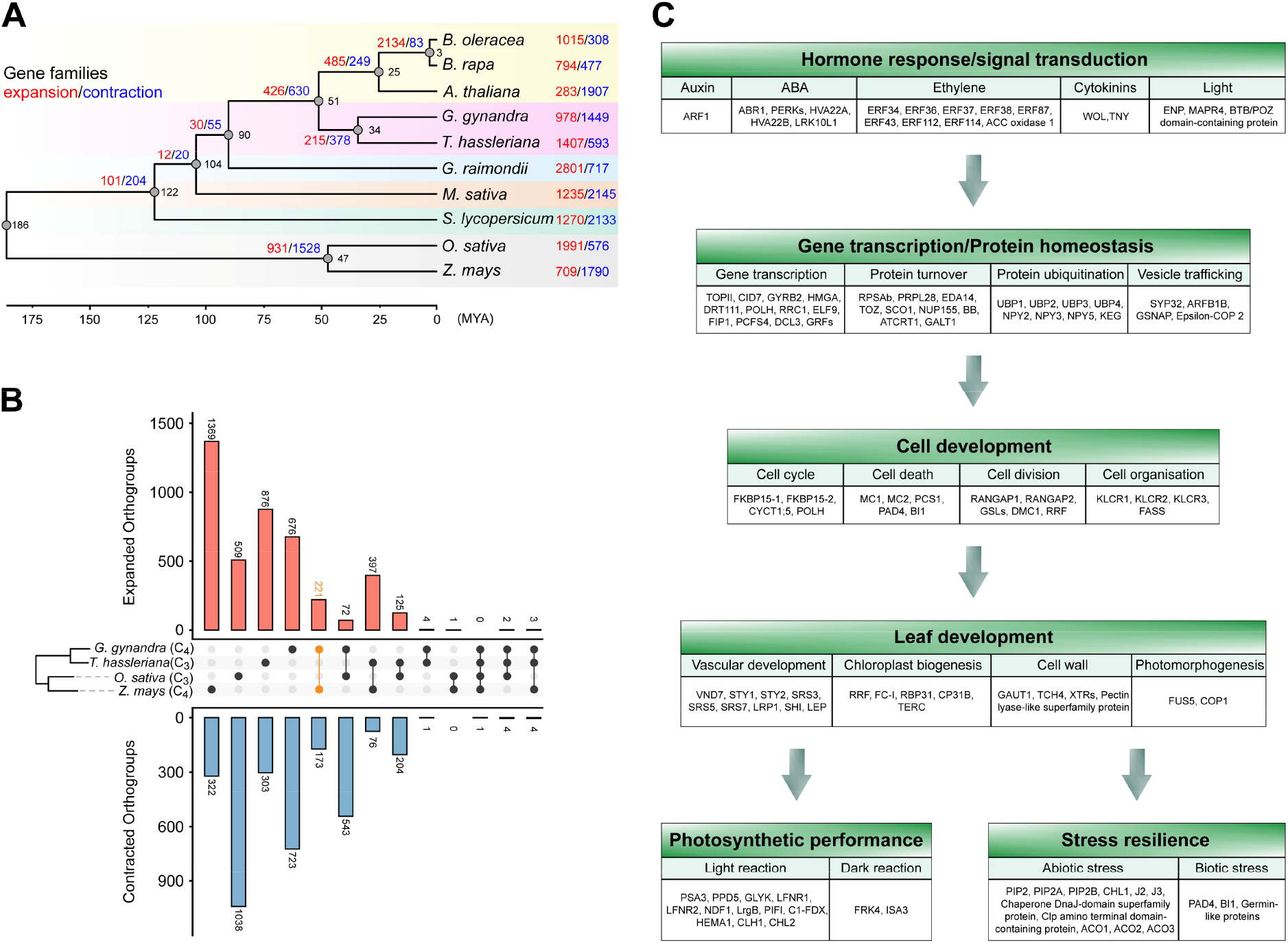
Analysis of gene family expansion and contraction in representative C_3_ and C_4_ plants. **(A)** The number of gene families that expanded (red) or contracted (blue) during evolution mapped to the species phylogenetic tree. The black number at each node denotes the divergence time between the two branches. The time scale (MYA) is shown at bottom. **(B)** Numbers of expanded and contracted orthogroups shared among the C_3_ and C_4_ lineages. The middle panels show phylogenetic tree and comparison groups of the four species, *G. gynandra* (C_4_), *T. hassleriana* (C_3_), *O. sativa* (C_3_) and *Z. mays* (C_4_). The upper and lower panels indicate the numbers of shared expanded and contracted orthogroups, respectively, with numbers noted for each bar. The number of expanded orthogroups shared between C_4_ plants *G. gynandra* and *Z. mays* is marked in orange (221). **(C)** Functional annotation of the expanded gene families common to *G. gynandra* and *Z. mays*. Arrows indicate biological processes potentially associated with the fitness advantages of C_4_ plants, including hormone response/signal transduction, gene transcription/protein homeostasis, cell development, leaf development, photosynthetic performance and stress resilience. The genes within each biological process are divided into different categories based on their functional annotations from MapMan.

### Gene expression patterns for high vein density and heat stress tolerance associated with maitenence of C_4_ photosynthesis in *G. gynandra*

The fitness advantages of C_4_ species are typically linked to alerations in leaf vein density or stress resistence during evolution. *G. gynandra* and *T. hassleriana* exhibited similar phenotypes regarding the morphology and leaf development (Figure 5A). However, *G. gynandra* possessed evidently denser veins (Figure 5B) and typical Kranz anatomy compared to *T. hassleriana* (Supplemental Figure S10). To identify potential discrepant features between these dicotyledonous C_3_ and C_4_ plants, their leaf samples from five developmental stages were collected under normal and heat stress conditions (Supplemental Figure S11). We found that *G. gynandra* had significantly higher vein density than *T. hassleriana* during early (S1), middle (S3) and mature (S5) stages (Figure 5C). Many of the well-known genes encoding vasculature developmental factors had differential expression patterns between the two species (Supplemental Figure S12). To dissect mechanisms underlying the differences in transcriptional regulation of the leaf vasculature, we constructed large-scale gene regulatory networks (GRN) based on RNA-seq time series datasets of *G. gynandra* and *T. hassleriana* using GENIE3, and candidate target genes of TFs were then predicted (Supplemental Figure S13). Among many of the TFs, the *Dof* family is established to be enriched in BSC and bind to AAAG motifs (Dai et al., 2022). Twenty-one and twenty-four hub genes from *Dof*-GRNs were identified for *G. gynandra* and *T. hassleriana*, respectively (Figure 5D). In addition to the well-documented *Dof* genes with important roles in vein development, such as *CDF2, CDF3, OBP2, OBP4, PEAR2, Dof2*.*4, HCA2* and *TMO6*, we also found some *Dof* genes with unknown functions, including *Vdof1, Dof1*.*4, Dof1*.*7, Dof1*.*8*, and *Dof2*.*2* (Supplemental Data Set S16). By analyzing of the previously deposited cell-specific transcriptomic data (Aubry et al., 2014), we found that both homologous genes of *Vdof1* in *G. gynandra* exhibited preferential expressions in BS cells (Supplemental Figure S14). The potential target genes of *G. gynandra Vdof1* were shown to be mainly enriched in pathways for chlorophyll metabolism, photosynthesis, and RNA and protein metabolism (Figure 5E). Overexpression of *Vdof1* conferred significantly higher photosynthetic efficiency, leaf vein density and heat resistance in *Arabidopsis* plants, whereas its corresponding mutants displayed the opposite effects (Supplemental Figure S15). Research on molecular mechanisms of *Vdof1* in regulating these C_4_ photosynthesis-associated processes is currently underway.

**Figure 5.**
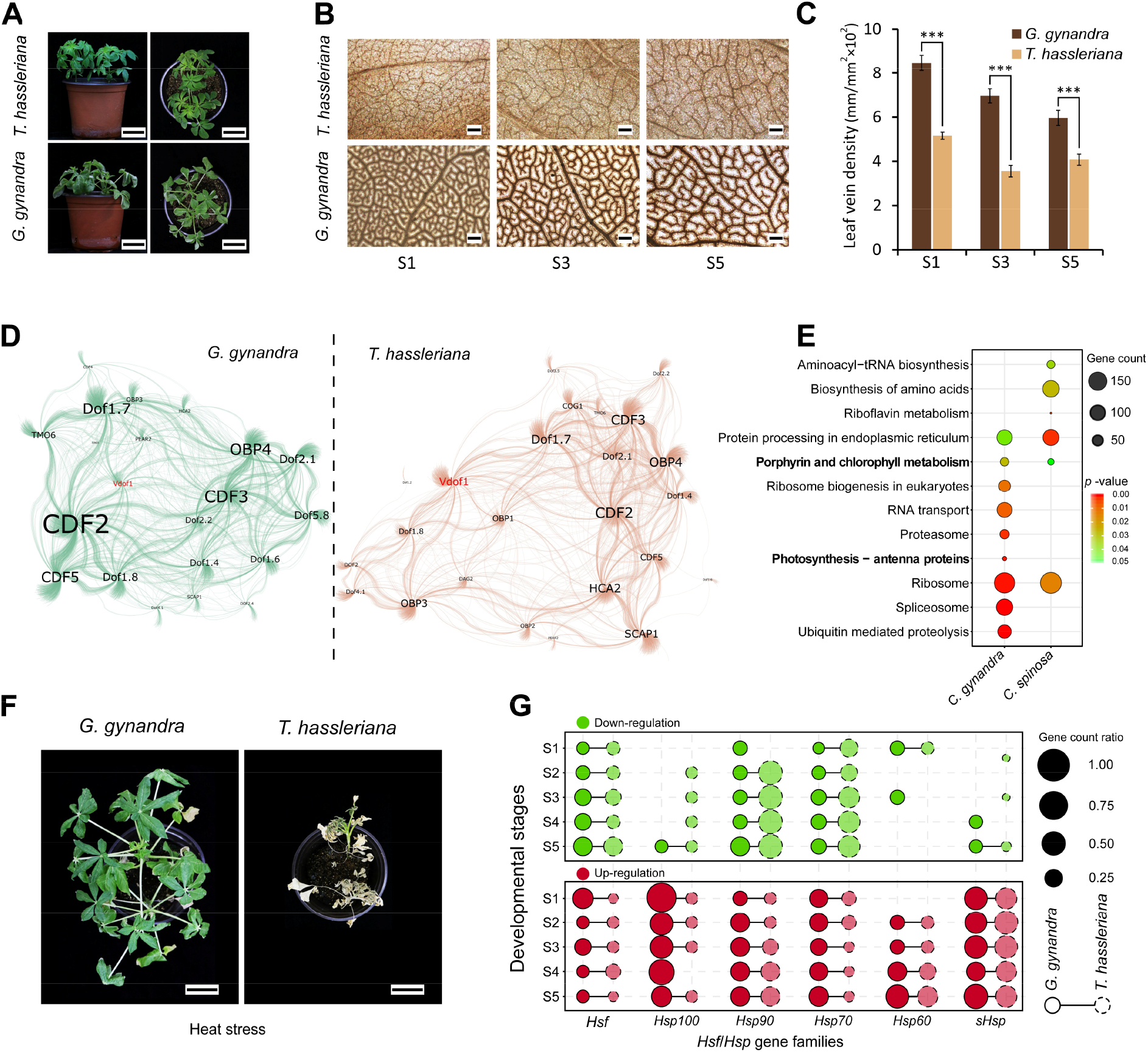
Higher leaf vein density and heat stress resistance in *G. gynandra* compared to *T. hassleriana* associated with gene expression modifications. **(A)** The growth phenotype of *G. gynandra* (lower panel) and *T. hassleriana* (upper panel) under normal conditions. The photographs are taken from 5-week old plants. The left and right panels are side and top views, respectively. Scale bar = 5 cm. **(B)** Overview of leaf vein patterns in *G. gynandra* (bottom) and *T. hassleriana* (top). Representative images from left to right illustrate vein density of the central region at leaf developmental stages from young to mature as numbered (S1, S3, S5). Scale bar = 200 μm. **(C)** Analysis of the leaf vein density between *G. gynandra* and *T. hassleriana*. Vein density (vein length/area) for each leaf was assessed for six positions at a central region bounded by the midvein over three developmental stages. Student’s *t* test (means ± SD; *n* = 3); \*\*\**P* < 0.001. **(D)** Networks of gene regulatory relationships between Dof transcription factors and their candidate target genes in *G. gynandra* and *T. hassleriana*. The gene regulatory networks (GRN) were constructed using GENIE3 and visualized with Gephi. The size of the labeled hub gene names is based on the degree of connecting nodes. The edges of *G. gynandra* (left) and *T. hassleriana* (right) GRNs are indicated in green and tangerine colors, respectively. The *Vdof1* gene in the two GRNs is highlighted in red. **(E)** KEGG pathway enrichment of the candidate target genes of the Vdof1 transcription factor in *G. gynandra* and *T. hassleriana*. The size of the circle indicates enriched gene numbers in each pathway, with the color of the circle indicating enrichment *p*-value. Pathways with *p*-value < 0.05 are shown. The photosynthesis-related pathways are marked in bold. **(F)** *G. gynandra* displays exceptional tolerance to continuous high temperature stress compared with *T. hassleriana*. 5-week-old plants were subjected to heat treatments in a controlled growth chamber under conditions of 45/35 °C (16/8h) and 60% relative humidity for 15 days. Scale bar = 5 cm. **(G)** The proportions of up- and down-regulated genes for *Hsf*/*Hsp* families after heat treatments in *G. gynandra* and *T. hassleriana*. The X-axis and Y-axis indicate *Hsf*/*Hsp* family names and different leaf developmental stages (from young to mature, S1 to S5), respectively. The size of the circle denotes the ratio of differentially expressed gene count relative to the total gene number of each *Hsf*/*Hsp* family. The lower (red) panel depicts the proportions of up-regulated genes, and the upper (green) panel shows the proportions of down-regulated genes.

We further examined the response of *G. gynandra* and *T. hassleriana* plants to heat stress treatment. Apparently, *G. gynandra* was much more tolerent to high teperature than *T. hassleriana* (Figure 5F). We then analyzed differential gene expression in the two species. A total of 16,138 and 18,492 differential expression genes (DEGs) were identified in *G. gynandra* and *T. hassleriana*, respectively (Supplemental Data Set S17). The heat stress response pathway was significantly up-regulated at all the stages in *G. gynandra* but not in *T. hassleriana* (Supplemental Figure S16). A set of *Hsf, Hsp100, Hsp90, Hsp70, Hsp60* and *sHsp* was identified in both species (Supplemental Data Set S18). However, 12–40% of *Hsf* genes in *G. gynandra* exhibited significantly up-regulated expressions after heat treatments, compared to 7–19% for *T. hassleriana*, depending on the leaf developmental stage (Figure 5G), and a larger number of *Hsp100, Hsp90, Hsp70*, and *Hsp60* genes were up-regulated in *G. gynandra*. Consistently, the photosynthesis pathway was over-represented among the heat-upregulated genes of *G. gynandra*, regardless of developmental stage (Supplemental Figure S16). The expression levels of a subset of C_4_ metablism-related genes were induced in *G. gynandra* under heat stress, such as *CA* (*GG03G120380*), *NAD-MDH* (*GG12G084140* and *GG09G089650*), *PEPC* copies, *PPDK, ALAAT* (*GG06G105780*), *ASPAT* (*GG07G009900*) and *NHD* copies, as were N assimilation pathway genes including *GS2* and *NiR* copies (Supplemental Figure S17). Carbon fixation, S assimilation and redox regulation pathways were largely unaffected, except for several *TPT* copies being up-regulated. While diffrential expression patterns were observed for photorespriration pathway genes between *G. gynandra* and *T. hassleriana*.

### Expression features and evolution of key genes involved in C_4_ photosynthesis and photorespiration

An illustration for NAD-ME subtype of C_4_ photosynthesis was shown (Figure 6A). Targeted analysis of the copy number of C_4_ cycle genes revealed that the ratio of gene copy number in *G. gynandra* vs *T. hassleriana* was largely higher than 2:3, which is the baseline expected ratio given their genomic relationship (Supplemental Figure S18). This suggests that *G. gynandra* likely retained more C_4_ genes after WGD, whereas *T. hassleriana* could have lost a subset of these genes after WGT (Supplemental Data Set S18). For instance, there were six copies of *NAD-MDH* in both species, nine and ten copies of *GAPDH* in *G. gynandra* and *T. hassleriana*, respectively, and even more copies of *GOGAT* (4:2) and *GDCP* (3:2) were found in *G. gynandra* compared to *T. hassleriana* (Supplemental Data Set S18). Furthermore, all the enzymes characterizing NAD-ME subtype were identified in *G. gynandra* (Figure 6B and Supplemental Data Set S18), and their gene expression in six different tissues were assessed. Transcript levels of the main candidate *NAD-ME* (*GG02G000360*) and *NAD-MDH* (*GG02G075940*) genes were over 1,406-fold and 7-fold higher in leaves than roots, respectively, whereas all the candidate *NADP-ME* and *PEP-CK* genes had drastically decreased expressions in leaves than in the other tissues. The expression levels of *Dit, NHD, BASS, TPT* and *AlaAT* genes were highest in leaves. These results indicate that C_4_ enzyme genes are preferentially expressed in photosynthetic tissues and that *G. gynandra* performs the NAD-ME subtype of carbon fixation. Futhermore, candidate homologous enzymes involved in C_4_ carbon fixation in *G. gynandra* were identified based on their preferential expression in photosynthetic tissues and phylogenetic analysis with known C_4_ genes (Supplemental Figure S19), which could be NAD-ME2 (GG02G000360), NAD-MDH/mMDH1 (GG02G075940 and GG02G075810), PEPC2 (GG14G004940, GG14G005080 and GG14G005010), PEPC-K/PPCK1 (GG05G088230), PPDK (GG13G000810), PPDK-RP/RP1 (GG12G002660), CA1 (GG15G098730 and GG10G000490) and CA4 (GG17G084960), ASPAT/ASP5 (GG07G010040), and ALAAT2 (GG06G105780).

**Figure 6.**
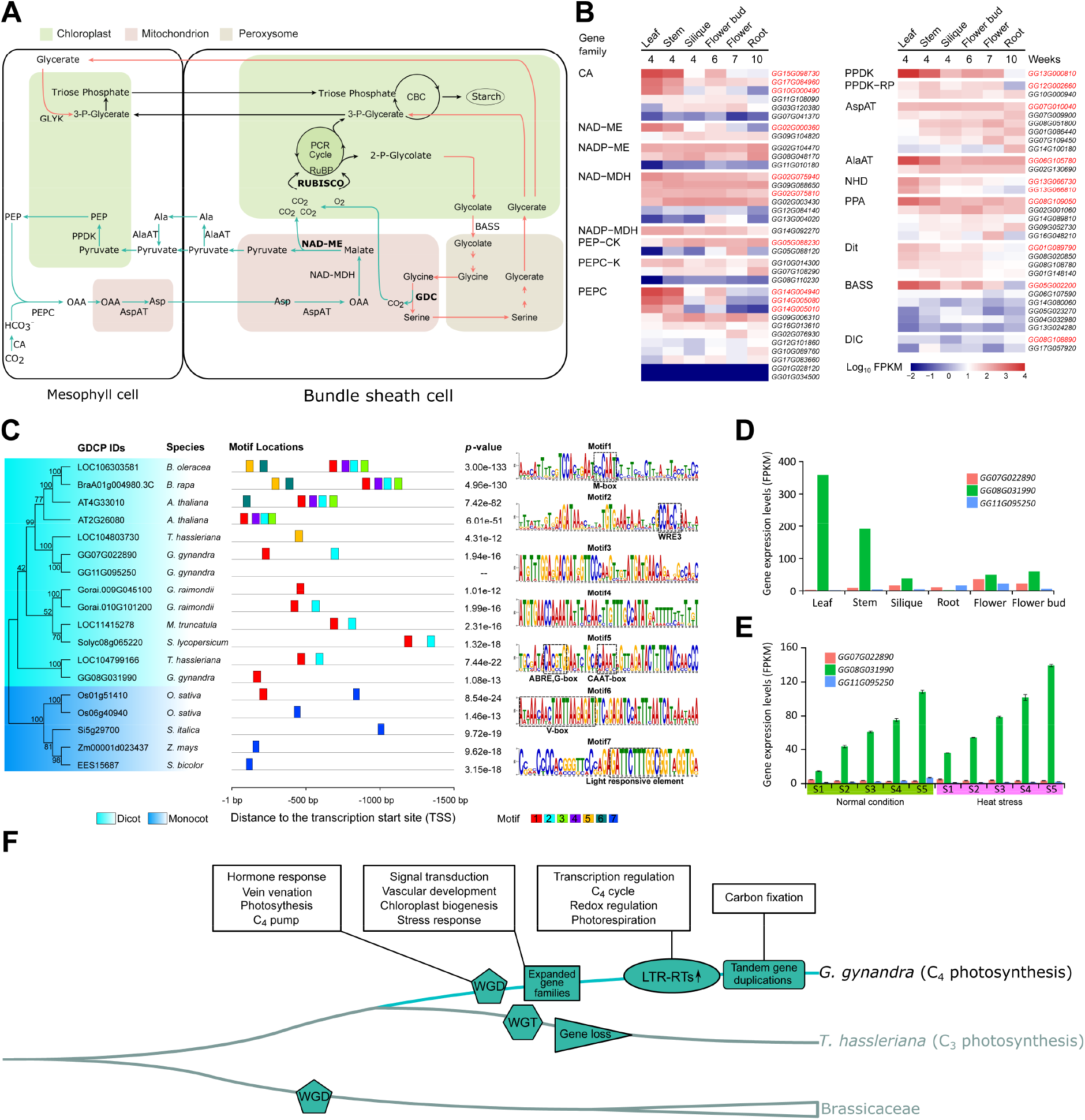
Evolution of C_4_ photosynthesis in *G. gynandra*. **(A)** Diagrammatic representation of main proteins and metabolic fluxes for the NAD-ME C_4_ photosynthetic subtype in *G. gynandra*. Aspartate (Asp) converted from oxaloacetate (OAA) by AspAT in the mitochondria of mesophyll (M) cells is the main metabolite transported from M cells to BS cells. Asp is converted to OAA and then to malate (Mal), which is decarboxylated by NAD dependent malate dehydrogenase (NAD-MDH); Mal is further decarboxylated by NAD-ME, releasing CO_2_ to the chloroplast of BS cells. The green arrows show the carbon dioxide (CO_2_) accumulation pathway of NAD-ME subtype. Red arrows mark the photorespiratory pathway. The three key enzymes, RUBISCO, NAD-ME and GDC, are in bold. Full names of metabolites and enzyme abbreviations are listed in Supplemental Data Set S21. PCR cycle, photosynthetic carbon reduction cycle; CBC, Calvin-Benson cycle. **(B)** Heatmap showing the expression pattern of key genes involved in C_4_ photosynthesis in photosynthetic and non-photosynthetic tissues of *G. gynandra*. The color denotes the expression level from low (blue) to high (red) expressed as log_10_FPKM. The ID of each gene copy in *G. gynandra* was indicated at right. The candidate C_4_ genes for *G. gynandra* are highlighted in red. **(C)** Phylogenetic tree and motifs in the promoter regions of *GDCP* genes in representative C_3_ and C_4_ plants. The left panel shows the phylogenetic tree of GDCPs, with light blue and dark blue backgrounds marking dicot and monocot species, respectively. The middle panel shows the type, location and *p*-values of conserved motifs in the upstream 1.5 kb regions from the transcription start site of each *GDCP* gene. The right panel shows the motif sequence and *cis*-acting regulatory elements highlighted by dashed boxes. The scale length (bp) and motif color key are shown at the bottom. **(D)** The expression patterns of the three *GDCP* genes in different tissues of *G. gynandra*. Unique among the paralogs, *GG08G031990* gene showed very high level of expression in the leaf or stem. **(E)** Analysis of expression levels of the three *GDCP* genes during *G. gynandra* leaf development under normal or heat stress conditions. The X-axis and Y-axis indicate gene expression level (FPKM) and leaf developmental stages (from young to mature, S1 to S5), respectively. Consistent with the results from Figure 6D, only *GG08G031990* was dominantly expressed in each stage analyzed, and its expression was even inducible by heat. **(F)** A conceptual model reconstructing hypothesized key steps in the evolution and maintenance of C_4_ photosynthesis in the spider plant. First, a recent whole-genome duplication (WGD, Gg-α) is the origin of C_4_ pathway-related genes in *G. gynandra*, including genes involved in the hormone response, vein venation, C_4_ metabolism, Calvin cycle, N and S assimilations and photorespiratory CO_2_ pump. Second, expansions of defined gene families enable the establishment of C_4_ features including signal transduction, vascular development, chloroplast biogenesis and stress response. Finally, more recent LTR-RT explosion and species-specific tandem duplications fine-tune the expression of C_4_ photosynthesis-associated genes, including those involved in the light reaction, C_4_ cycle, redox regulation and carbon fixation. While these steps can overlap and may not have taken place independently. The much higher leaf vein density and heat stress tolerance in *G. gynandra* compared to *T. hassleriana* are linked with shifted patterns of gene expression. Despite *T. hassleriana* underwent a whole-genome triplication (WGT, Th-α) event later than Gg-α, subsequent massive gene loss likely occurred, including missing the duplicated copies of certain C_4_ metabolism genes. Besides, contrary to the photorespiration pathway, the overall expression levels of the C_4_ pathway-related genes in *T. hassleriana* are markedly lower than in *G. gynandra*, which might have also constrained its evolution of C_4_ photosynthesis.

The restriction of Gly decarboxylase (GDC) to BSC results in formation of the photorespiratory CO_2_ pump that is an essential step during C_3_-to-C_4_ transition. This promoted us to examine how P-protein of GDC (GDCP) and its promoter elements are altered in C_3_, C_4_ dicot and C_4_ monocot species (Figure 6C). Interestingly, three copies of *GDCP* gene were detected in *G. gynandra*, while a single copy was present in C_4_ monocots (*S. bicolor, Sorghum italica* and *Z. mays*) and 1–2 copies were in C_3_ plants. A total of seven conserved motifs (motif 1 to 7) were identified in promoter regions of these *GDCP* genes. Motif 1 containes an element (M-box) that has reported to drive expression in mesophyll cells (MC), with motif 6 harboring a regulatory element (V-box) being required for vasculature expression (Adwy et al., 2015, 2019). Unexpectedly, M-box, but rather V-box, was present in the promoters of two *GDCP* genes (*GG08G031990* and *GG07G022890*) in *G. gynandra*, and no known motifs could be detected in the other one *GG11G095250*. It should be metioned that neither M-box nor V-box was found in the *GDCP* of C_4_ monocots. Notably, a single *GDCP* (*GG08G031990*) was the most abundantly expressed in *G. gynandra*, with the highest levels in leaf tissue, while the other two duplicates were expressed at extremely low levels in all tissues analyzed (Figure 6D). This paralog was uniquely increased along leaf developmental gradients and could be further dramatically induced by heat stress (Figure 6E). Based on our data, we proposed a hypothetical model for the evolution and maintenance of C_4_ photosynthesis in the spider plant (Figure 6F).

## DISCUSSION

*G. gynandra* has been an instrumental organism for addressing evolutionary questions, often as a pair with *T. hassleriana* (Hoang et al., 2022). This plant is valuable for the study of its nutritional and health-promoting properties, as well as molecular mechanisms absent in C_3_ model organisms such as the dicot *Arabidopsis* and the monocot rice, and represents an emerging model plant to extensively investigate C_4_ biology (Brown et al., 2005; Külahoglu et al., 2014; Bayat et al., 2018). However, the lack of a high-quality chromosome-scale genome of *G. gynandra* has hampered its wide adoption in basic and applied research. Here we report a reference-grade genome of the diploid *G. gynandra* integrating long reads Nanopore sequencing and Hi-C technologies to overcome the challenge inherent in highly repetitive sequences (Figures 1 and 2). With a contig N50 of 11.43 Mb, scaffold N50 of 51.02 Mb and LAI of 16.46, the assembly metrics indicate high accuracy and completeness (Table 1). Genomics combined with transcriptomic analysis confirmed that *G. gynandra* is the typical NAD-ME subtype of C_4_ plant (Figure 6). Given that an *Agrobacterium tumefaciens*-mediated transformation system with high efficiency has also been established for this elite crop (Newell et al., 2010), *G. gynandra* is now suitable as a C_4_ model system in dicotyledonous plants, particularly for studies in NAD-ME subtype of C_4_ species, such as the perennial polyploid bioenergy crop switchgrass (*Panicum virgatum*).

The evolution of plant C_4_ photosynthesis represents an unique example of convergent evolution (Heyduk et al., 2019; Wang et al., 2009; Sage, 2004). With the availability of the annotated genomes for *G. gynandra* and *T. hassleriana*, direct comparison between these C_4_ and C_3_ species presents an excellent model to unravel why C_4_ photosynthesis evolved in only one of the two plants. Instead of sharing a Th-α event (van den Bergh et al., 2014), these two species experienced independent recent WGD (Gg-α) and WGT (Th-α) events, respectively (Figure 3). However, massive gene loss may have happened in C_3_ *T. hassleriana* after Th-α (Supplemental Figure S18 and Supplemental Data Set S19), which resulted in retention of only 27,396 predicted genes, making *T. hassleriana* the gene-poorest species of those analyzed here (Figure 1D). Of note, the copy numbers of multiple C_4_ pathway-related genes are higher in *G. gynandra* relative to *T. hassleriana* (Supplemental Data Set S18), including those involved in vasculature development (*PIN1, SCR* and *HB-8*), C_4_ metabolism (*ALAAT* and *NHD*), Calvin-Benson cycle (*RbcSSU*), N assimilation (*FdGOGAT*), S assimilation (*APR* and *SR*), and photorespiration (*GDCP* and *SGAT*), concomitant with markedly reduced expressions of these vasculature development and C_4_ metabolism genes in *T. hassleriana* (Supplemental Figure S20). Moreover, the Gg-α in *G. gynandra* contributed to the expansion of some gene families shared with the C_4_ monocot *Z. mays*, which function in various C_4_ metabolism-associated processes from signaling pathways to photosynthetic performance and stress resistance (Figure 4). This is supportive of recent studies showing that WGD led to multiple aspects of the evolution of C_4_ photosynthesis in *G. gynandra* and grasses (Huang et al., 2021, 2022). Almost all of the well-documented vein development-associated genes in *G. gynandra* have retained their WGD-derived duplicates (Supplemental Data Set S20) (Ohashi-Ito and Fukuda, 2010; Huang et al., 2021; Liu et al., 2022). Importantly, some of these paralogous genes that positively control vein density became significantly up-regulated in *G. gynandra* compared to *T. hassleriana*, with those negative ones being dramatically down-regulated (Supplemental Figure S12) (Huang et al., 2017; Aubry et al., 2014; Liu et al., 2020), which may have conferred the much higher leaf vein density seen in *G. gynandra* (Figure 5, B and C). Peculiarly, *G. gynandra* has retained the paralogous genes essential for building the photorespiratory CO_2_ pump in BS cells, such as *ALAAT, ASPAT, GOGAT* and *GDCP* (Supplemental Data Sets S18 and S20) (Huang et al., 2021), whose formation is hypothesized to be the primary driver of C_4_ evolution (Mallmann et al., 2014; Sage et al., 2012). Although the overall level of expression for photorespiration pathway genes in *G. gynandra* is down-regulated, photorespiration is not and cannot be lost, as it has crucial functions in plants. These data suggest that *G. gynandra* underwent the C_3_-C_4_ intermediate stage required to establish the C_4_ pathway. *G. gynandra* harbors three *GDCP* gene copies; however, only one is highly expressed in BS cells, which exhibits gradually increased expression during leaf development (Figure 6, C–E). It is speculated that the inactivated gene copy will eventually be lost (Schulze et al., 2016; Mercado and Studer, 2022). Consistently, a single *GDCP* is active in monocot C_4_ species (maize, foxtail millet and sorghum) and in the dicot C_3_-C_4_ intermediate species *Flaveria* and *Moricandia* (Figure 6C) (Schulze et al., 2013; Schlüter et al., 2017). These imply that the C_4_ trait in *G. gynandra* could be still evolving. Additionally, genomic analysis showed that the promoter regions of the *GDCP* genes in C_3_, C_4_ dicots and C_4_ monocots contained diverse *cis*-active element motifs (Figure 6C). While the V-box element has been demonstrated to orchestrate the expression of *GDCP* in the vasculature of *Moricandia* (Adwy et al., 2019, 2015), it is not found in *G. gynandra* or C_4_ monocots, implying that varied modes might have evolved to regulate cell-specific *GDCP* expression not only between C_3_ and C_4_ species but also among C_4_ plants.

Heat, intense light and/or drought stress were proposed to drive C_4_ origination and evolution during periods of declining atmospheric CO_2_, suggesting the role of adaptive evolution in C_4_ pathway formation (Blätke and Bräutigam, 2019; Sage, 2004). Not surprising, C_4_ photosynthesis has better performance than the ancestral C_3_ state under stressful environments. This will be increasingly important under the projected conditions of long-term climate change and periodic environmental extremes currently threatening the global food supply and bioenergy security. *G. gynandra* displays much higher tolerance to extreme temperature than *T. hassleriana*, and this phenotype is likely associated with retention of a subset of WGD-duplicated genes in *G. gynandra* that encode HSFs and HSPs (Figure 5F and Supplemental Data Set S20). Although their expressions can be induced under heat stress in both species, many more of these families are up-regulated in *G. gynandra* compared to *T. hassleriana* (Figure 5G). Given that C_4_ photosynthesis in *G. gynandra* evolved after WGD (van den Bergh et al., 2014), this provides additional evidence on the critical role of Gg-α in facilitating C_4_ evolution.

Aside from WGD event and the expanded gene families, recent species-specific tandem duplications after Gg-α may also be relevant to maintenance of the C_4_ pathway in *G. gynandra*, exemplified by duplication of multiple key enzymes involved in carbon fixation including NAD-MDH/mMDH1 and PEPC2 of C_4_ cycle (Figure 3, E–G). Remarkably, our data unveiled a previously undescribed role for genomic transposon expansion in the evolution of C_4_ photosynthesis. The higher frequency of LTR-RT insertion in promoter regions of *G. gynandra* contributes to higher ratio of up-regulated than down-regulated genes when compared to *T. hassleriana*, independent of leaf developmental stages. Of importance, the up-regulated genes function in various pathways involved in C_4_ photosynthesis, including but not limited to, C_4_ cycle, light reaction, cyclic electron flow, Calvin cycle, redox regulation, auxin signaling, vein patterning and stress response, whereas LTR-RT-inserted genes involved in photorespiration exhibit lower expressions (Figure 2). Besides, a small portion of C_4_ homologous genes that contain less and more LTR-RT insertions, whose expressions are up-regulated and down-regulated, respetively (Supplemental Figure S6C). Therefore, it is tempting to postulate that LTR-RT activity might have promoted the C_3_-to-C_4_ evolution via synergistic genomic and transcriptomic alterations. Consistent to the recent reports demonstrating that modifications in *cis-*regulatory and non-coding regions are primary determinants of the distinct transcriptome blueprint in BSC or MC of C_4_ plants (Tu et al., 2020; Dickinson et al., 2020; Dai et al., 2022), this extensive LTR-RT insertion frequency in promoter regions appears to be a underappreciated route through which cell-preferential gene expression could be achieved. Future molecular characterization of regulatory regions containing LTR-RTs will be required to test this hypothesis. Altogether, our present study suggests a three-step scenario contributing to the evolution of C_4_ photosynthesis in the *Cleome* genus and that innovation of this novel biochemical pathway in *G. gynandra* may be dual consequences of duplicated gene retention and functional changes in existing genes (Figure 6F).

In summary, we have generated a highly continuous and accurate genome assembly of the C_4_ model crop *G. gynandra*. Systematic comparison with the closely related C_3_ species *T. hassleriana* is useful for identification of previously uncharacterized genes involved in C_4_ photosynthesis-related processes. Moreover, we present evidence that LTR-RT bursts have functioned in driving genome size increase and may have been important for C_4_ evolution as well. WGD, gene family expansion and tandem duplication events have also facilitated the evolution of C_4_ photosynthesis in *G. gynandra*. We found corresponding transcriptional changes of gene expression features associated with C_4_ pathway, photorespiration, vein venation and heat stress response for boosting photosynthetic efficiency. These findings shed light on the commonalities and differences in the evolution of C_4_ photosynthesis, supporting the existence of numerous independent evolutionary trajectories to C_4_. The genomic and transcriptomic data generated in this study provide valuable new resources for further dissecting the genetic basis underlying the transition from C_3_ to C_4_ photosynthesis as well as the exceptional nutritional and medicinal traits of this species. *G. gynandra* is thus now a promising model species to accelerate both basic and applied research, in C_4_ biology and beyond.

## MATERIALS AND METHODS

### Plant materials and growth conditions

C_4_ *Gynandropsis gynandra* Linn. and C_3_ *Tarenaya hassleriana* (Purple Queen) species were grown in plastic pots with Pro-Mix BX soil, in a controlled environment room with a photoperiod of 16h/8h, a light intensity of 200 µmol m^-2^ s^-1^, 25/18 °C (day/night), and a humidity of 50–60%. The leaves (4 weeks), stems (4 weeks), roots (4 weeks), flower buds (6 weeks), flowers (7 weeks) and siliques (10 weeks) of plants were harvested and immediately frozen in liquid nitrogen.

### Heat treatment

5-week-old plants of both species were transferred to a growth chamber (PercivalE-41, USA) using the conditions above, except with the day/night temperature set to 45/35 °C, which is 10–15 °C above optimum range of germination for *G. gynandra* and is considered as heat stress (Motsa et al., 2015). Leaves at different developmental stages were sampled from plants at 10:00 am before and after heat treatment. Meanwhile, leaf samples at the equivalent stages under normal growth conditions were collected as controls (Supplemental Figure S11). Stages 1 to 5 (S1 to S5) were leaves from young to mature, with Stage 1 as the youngest (1.5 cm in length). All experiments were performed in triplicate.

### RNA-Seq analysis

For RNA-Seq experiments, total RNAs were extracted and purified from the collected leaf samples above using Trizol RNA extraction kit (Invitrogen, Carlsbad, CA, USA). The mRNA was enriched from approximately 50 ng of high-quality total RNA with NEXTflex™ Poly(A) Beads (Bioo Scientific, Austin, TX, USA), and used to produce RNA-seq libraries with the NEBNext Ultra RNA Library Prep Kit for Illumina (New England Biolabs, Ipswich, MA, USA) according to the manufacturer’s protocol. The RNA-Seq libraries were sequenced with Illumina HiSeq4000 (Illumina, San Diego, CA, USA) under PE150 mode. The raw reads of RNA-seq were filtered using Trimmomatic (version 0.40) with default parameters (Bolger et al., 2014). The remaining clean reads were mapped to the genome using HISAT2 (version 2.1.0) (Kim et al., 2019). The reads count per gene was calculated with HTSeq (version 0.9.172) (Anders et al., 2015). The differentially expressed genes (DEGs) were identified using the DEseq2 package with a significance threshold of *q* value < 0.01 and |log_2_(fold change)| > 1 (Love et al., 2014).

### Iso-Seq analysis

Total RNAs extracted from different tissues collected above were combined to generate a full RNA sample of *G. gynandra*. The mRNA was enriched from total RNA using a magnetic d(T) bead binding procedure, and then was transcribed to cDNA using the Clontech SMARTer PCR cDNA Synthesis Kit (Clontech, Mountain View, CA, USA) following the manufacturer’s instructions. The amplified cDNA fragments were size-selected using a BluePippin Size Selection System (Sage Science, Massachusetts, United States) with a bin of >4 kb. The amplified and size-selected cDNA products were used to generate SMRT Bell libraries according to the Iso-Seq protocol (P/N100-377-100-05 and P/N100-377-100-04). The libraries were prepared for sequencing by annealing a sequencing primer and binding polymerase to the SMRT bell templates using the DNA/Polymerase Binding Kit (Pacific Biosciences, Menlo Park, CA, USA). One SMRT cell was sequenced on the PacBio Sequel instrument (Pacific Biosciences, Menlo Park, CA, USA). The high-quality, full-length and consistent transcript sequences were obtained from long reads data using the PacBio Iso-Seq3 pipeline.

### Genome sequencing

Nanopore, Illumina HiSeq and Hi-C were used for sequencing the complete genome of *G. gynandra*. For Illumina sequencing library construction, the genomic DNA of *G. gynandra* was extracted from young leaves using a Qiagen DNA purification kit (Qiagen, Darmstadt, Germany). The integrity of DNA was assessed by agarose gel electrophoresis, and the purity and concentration were determined on NanoDrop 2000 spectrophotometer (Thermo Fisher Scientific, MA, USA) and Qubit Fluorometer (Invitrogen, Carlsbad, CA, USA), respectively. The short-read genomic sequencing libraries were constructed using the MGIEasy FS DNA Library Prep Set (Item No.1000006988) with 270 bp fragment size according to the manufacturer’s instructions, and sequenced on an Illumina Hiseq4000 platform (Illumina, San Diego, CA) to generate paired-end (PE) reads of 150 bp. For Nanopore sequencing library preparation, the high-quality genomic DNA of *G. gynandra* was extracted from young leaves using the CTAB method. Approximately 15 μg of genomic DNA was used to collect larger DNA fragments (>20 kb) with the BluePippin Size-Selection system (Sage science, Beverly, MA, USA). The obtained genomic DNA was processed using ONT Ligation Sequencing Kit (SQK-LSK109; Oxford Nanopore Technologies, Oxford, United Kingdom) according to the manufacturer’s instructions. Briefly, the genomic DNA was end-repaired and dA-tailed using a NEBNext UltraII End Prep Reaction Module (New England Biolabs, Ipswich, MA, USA). The sequencing adaptors were ligated using NEBNext UltraII Ligation module (New England Biolabs, Ipswich, MA, USA). Libraries were purified using AMpure XP beads (Beckman Coulter, California, USA) and short fragment buffer (SFB). Then, Nanopore libraries were added into a single ONT MinION R9.4 flowcell (FLO-MIN106) and sequenced on the platform of PromethION (Oxford Nanopore Technologies, UK). The ONT MinKNOW software (https://github.com/nanoporetech/minknow_lims_interface.git) acquired raw sequence data with live basecalling by ONT Guppy (https://nanoporetech.com/). For Hi-C sequencing, the fresh leaf tissues of *G. gynandra* were fixed (cross-linking) with formaldehyde. The cross-linked chromatin was extracted and digested by *Mbo*I restriction enzyme, the 5’ overhangs filled in with biotinylated nucleotides, and blunt-end proximity-ligated to generate circular molecules. Subsequently, the circular DNA molecules were purified from protein, and sheared to ~350 bp mean fragment size, and then enriched by biotin pull down. The sequencing libraries were generated using NEBNext Ultra enzymes (New England Biolabs, Ipswich, MA, USA) and Illumina-compatible adapters (Illumina, San Diego, CA, USA). The Hi-C libraries were processed to paired-end sequencing on the Illumina Hiseq 4000 (Illumina, San Diego, CA, USA) platform with read length of 150 bp.

### Genome size estimation

The genome size of *G. gynandra* was estimated using the *k*-mer analysis. Raw reads from Illumina sequencing were subjected to SOAPnuke (https://github.com/BGI-flexlab/SOAPnuke) for base quality control. The genome size, abundance of repetitive elements and heterozygosity were estimated based on the *k*-mer frequencies generated from the short reads using GenomeScope (https://github.com/schatzlab/genomescope).

### *De novo* genome assembly

The ONT long reads were corrected and assembled using CANU (version 1.7.1) with the parameters (corOutCoverage=50, saveOverlaps=TRUE, ovsMemory=64, minMemory=30G, batMemory=200G, minOverlapLength=700, minReadLength=1000) (Koren et al., 2017). The resulting contigs were polished with both long reads and short reads using three rounds of Racon (version 1.4.13) with default parameters (Vaser et al., 2017). The Illumina short reads were mapped onto the polished assembly with BWA-MEM (https://github.com/bwa-mem2/bwa-mem2). Based on the alignment, error correction was conducted using Pilon (version 1.23) (Walker et al., 2014). To scaffold the assembled contigs, the ONT-based polished contigs were anchored into a chromosome-scale assembly using a Hi-C proximity-based assembly approach. Illumina reads from the Hi-C library were processed with SOAPnuke to remove adaptor and low-quality sequences. The clean Hi-C read pairs were used as input for the Juicer (version 1.6) and 3d-DNA Hi-C analysis and scaffolding pipelines (version 180922) (Dudchenko et al., 2017; Durand et al., 2016). Valid interaction pairs were mapped onto the polished contigs and anchored to the pseudo-chromosomes using 3d-DNA pipeline with default parameters. The Hi-C interaction matrix was visualized using Juicebox Assembly Tools, and mis-assemblies and mis-joins were manually corrected based on neighboring interactions to generate the final pseudo-chromosome-length genome assembly.

### Genome assembly quality assessment

Three assessment strategies including BUSCO alignment, transcriptome alignment and LTR assembly index, were used to evaluate the quality and completeness of the *G. gynandra* genome assembly. In brief, the completeness of the genome assembly was evaluated using the BUSCO pipeline based on the datasets of embryophyta_odb10. The coverage and base-level accuracy of the genome assembly were assessed by aligning transcriptome reads to the *G. gynandra* assembly using HISAT2 with default parameters. LAI score of the *G. gynandra* genome assembly was calculated using LTR_retriever (version 2.9.0) with default parameters (Ou and Jiang, 2018).

### Genome annotation

The *G. gynandra* genome was annotated including annotations of repeat sequences, protein-coding genes and RNA genes. First, we adopted two complementary methods (one homology-based and the other *de novo*-based) to predict repeat sequences. RepeatMasker (version 4.0.7) and RepeatProteinMask (http://www.repeatmasker.org) were used to discover and classify repetitive sequences with the homology-based library generated from Repbase (www.girinst.org/repbase) (Bedell et al., 2000). LTR_FINDER (version 1.06) and RepeatModeler (http://www.repeatmasker.org/RepeatModeler) were utilized to build the *de novo*-based library (Xu and Wang, 2007). TRF (version 4.09) was employed to annotate tandem repeats with default parameters (Benson, 1999). Full length LTR-RTs were identified using LTR_retriever. Second, we deployed a strategy combining *ab initio*, homology-based and transcriptome-based methods for gene structure annotation. We used both Augustus (version 3.3.2) and SNAP (version 1.0.4) to perform *ab initio* predictions with self-trained prediction models (Stanke et al., 2004; Korf, 2004). We aligned the protein sequences from *A. thaliana, B. oleracea, B. rapa, T. hassleriana, G. raimondii, M. Sativa, S. lycopersicum, O. sativa* and *Z. mays* to the repeat-masked genome of *G. gynandra*, and then parsed the resultant alignments by GeneWise (https://www.ebi.ac.uk/~birney/wise2/) to achieve homolog predictions. We mapped the transcripts generated from both PacBio Iso-Seq and Illumina RNA-Seq to the genome with PASA (version r20140407) to conduct transcriptome-based predictions (Haas et al., 2008). Finally, we combined all the evidences to finalize the consensus gene models using Maker (version 2.31.9) (Cantarel et al., 2008). Third, we performed RNA genes annotation by tRNAscan-SE (version 2.0) for tRNA (Lowe and Chan, 2016), RNAmmer (version 1.2) for rRNA (Lagesen et al., 2007), and INFERNAL (version 1.1.2) for miRNA and snRNA (Nawrocki and Eddy, 2013). In addition, we annotated transcription factor encoding genes using iTAK (version 1.7) (Zheng et al., 2016). We generated functional assignments of protein coding genes by performing BLAST searches against five public protein databases including the NCBI non-redundant (nr) database, SwissProt database, Clusters of Orthologous Genes (COG) database, KEGG database and GO database. Pfam domains of genes were identified using InterProScan.

### Phylogenomic evolution analysis

Orthology prediction for ten species including eight eudicots (*A. thaliana, B. oleracea, B. rapa, G. gynandra, T. hassleriana, G. raimondii, M. Sativa* and *S. lycopersicum*), two monocots (*O. sativa* and *Z. mays*) was performed using OrthoFinder (version 2.2.6) package with default parameters (Emms and Kelly, 2019). Single-copy orthologous genes were extracted from the clustering results, and were used to reconstruct the phylogenetic tree. In brief, the proteins of single-copy gene families were aligned by MUSCLE (version 3.8.31) (Edgar, 2004). The alignments were finally joined into a super gene matrix for the species phylogenetic tree construction using RAxML (version 8.2.12) (Stamatakis, 2014) with the JTT+I+GAMMA model and 1,000 bootstrap replicates. The best suitable evolution model for phylogeny construction was evaluated using jModelTest (version 2.1.10) (Darriba et al., 2012). Time estimation among species was performed with r8s (version 1.81) (Sanderson, 2003). The divergence times of *B. oleracea* and *B. rapa* (2.02–3.21 MYA), *Z. mays* and *O. sativa* (42–52 MYA) and *A. thaliana* and *O. sativa* (115–308 MYA) obtained from TimeTree (http://timetree.org) were used for calibration. Syntenic blocks and gene duplications were identified within the *G. gynandra* genome or between the *Cleome* plants and other species using MCScanX (version 0.8) with the parameters (-s 5, -m 5) (Wang et al., 2012). The synonymous mutation rate values for gene pairs within syntenic blocks were calculated using the PAML (version 4.9) with yn00 and NG model (Yang, 2007). Fourfold degenerative transversion (4DTv) rates were calculated by aligning all orthologous or paralogous gene pairs using an in-house Perl script. We identified the WGD event within the *G. gynandra* genome using the approach from PGDD (Lee et al., 2013). CAFE (version 4.2.1) was used to identify the gene families with rapid expansion or contraction based on the species phylogenetic tree and divergence time (De Bie et al., 2006). The species-specific gene families were determined according to the presence and absence of genes for specific species.

### Analysis of leaf anatomy

Leaf samples were rinsed twice in water and placed in 70% ethyl alcohol for 3–4 days. The 70% ethyl alcohol was changed every eight hours until the leaves became colorless and transparent. Leaves were rinsed again with water and mounted in 66% glycerol on slides for observation. Images of the cleared leaves were taken with an OLYMPUS BX43 microscope system equipped with an OLYMPUS DP74 camera at 2× magnification (Olympus, Tokyo, Japan). Vein length and area size were assessed for six sites at a central region of each leaf bounded by the midvein using phenoVein tools developed from MeVisLab (www.mevislab.de) (Bühler et al., 2015). Vein density (Vd) was calculated as the total length of all veins within the region divided by the area of the region, which was expressed in mm mm^-2^. Measurements were conducted with three independent leaves of each stage per species.

### Phylogenetic and cis-acting regulatory elements analysis

The protein sequences of *GDCP* genes were used to perform multiple sequence alignment using MUSCLE. Phylogenetic tree was inferred using the Neighbor-Joining (NJ) method available in MEGA7 (version 7.0.26) (Kumar et al., 2016). The robustness of each node in the tree was determined using 1000 bootstrap replicates. The upstream 1.5 kb regions from transcription start site (TSS) of each *GDCP* gene were extracted from genome sequence of each species, and used to identify conserved motifs using MEME, and *cis*-acting regulatory elements by PlantCARE server (http://bioinformatics.psb.ugent.be/webtools/plantcare/html).

### Accession Numbers

All sequence information has been uploaded to Genome Sequence Archive (https://bigd.big.ac.cn/gsa). Genomic data: *Gynandropsis gynandra*-PRJCA008306; Transcriptomic data: *Gynandropsis gynandra*-PRJCA008307; *Tarenaya hassleriana*-PRJCA008308. The chromosome-scale annotated genome assembly of *Gynandropsis gynandra* is available at the Genome Warehouse in the National Genomics Data Center (https://ngdc.cncb.ac.cn) under accession number GWHBHRJ00000000.

## Supplemental Files

Supplemental Figure S1. Statistics for the frequencies and depths of 15 *k*-mers in the *G. gynandra* genome.

Supplemental Figure S2. Overview of the processing pipeline for assembling the *G. gynandra* genome.

Supplemental Figure S3. Length distribution of ONT sequencing reads.

Supplemental Figure S4. KEGG functional enrichment analysis of gene families specific to *G. gynandra*.

Supplemental Figure S5. The density distribution of intact LTRs, *Gypsy* and *Copia* in each chromosome of *G. gynandra*.

Supplemental Figure S6. Genomic distribution of LTR-RTs and their impacts on expression patterns of the C_4_ pathway-associated genes in *G. gynandra* compared to *T. hassleriana*.

Supplemental Figure S7. Comparative genomic analysis of the WGD event in *G. gynandra*.

Supplemental Figure S8. Functional enrichment of WGD-derived genes and expression analysis of TD-derived genes in *G. gynandra*.

Supplemental Figure S9. MapMan-Bin enrichment analysis of 221 expanded gene families common to *G. gynandra* and *Z. mays*.

Supplemental Figure S10. Representative anatomy of the leaf cross-section of *G. gynandra* and *T. hassleriana*.

Supplemental Figure S11. The morphology of leaves at different developmental stages for *G. gynandra* and *T. hassleriana*.

Supplemental Figure S12. The expression patterns of leaf and vasculature-related regulatory genes along leaf development gradients for *G. gynandra* and *T. hassleriana*.

Supplemental Figure S13. The gene regulatory networks (GRNs) between transcription factors and their candidate target genes in *G. gynandra* or *T. hassleriana*.

Supplemental Figure S14. Enrichment of *Vdof1* gene in leaf BS cells.

Supplemental Figure S15. Overexpression of *Vdof1* gene led to enhanced photosynthetic capacity, leaf vein density and tolerance to heat stress.

Supplemental Figure S16. MapMan-Bin enrichment analysis of DEGs at different stages of leaves from *G. gynandra* and *T. hassleriana* subjected to heat stress.

Supplemental Figure S17. Heatmap showing expression patterns of C_4_ photosynthesis-related pathway genes under heat stress in *G. gynandra* and *T. hassleriana*.

Supplemental Figure S18. Copy number ratio of genes for C_4_ photosynthesis-related pathways in *G. gynandra* as compared to *T. hassleriana*.

Supplemental Figure S19. Phylogenetic trees of C_4_ photosynthesis-related genes.

Supplemental Figure S20. The expression levels of C_4_ pathway-related genes with lower copy numbers in *T. hassleriana* than in *G. gynandra* along leaf development gradients.

Supplemental Data Set S1. Estimate of *G. gynandra* genome size.

Supplemental Data Set S2. The evaluation of ONT sequencing data for *G. gynandra*.

Supplemental Data Set S3. Statistics of Hi-C data for *G. gynandra*.

Supplemental Data Set S4. Summary of *G. gynandra* genome assembly.

Supplemental Data Set S5. Summary of *G. gynandra* seventeen pseudo-chromosomes. Supplemental Data Set S6. BUSCO analysis of *G. gynandra* genome assembly.

Supplemental Data Set S7. RNA-seq data mapping summary.

Supplemental Data Set S8. Functional annotation of the predicted genes in the assembly of *G. gynandra* genome.

Supplemental Data Set S9. Statistics of the annotated non-coding RNAs.

Supplemental Data Set S10. Gene numbers of transcription factor family among *G. gynandra* and nine other species.

Supplemental Data Set S11. Numbers of gene families among *G. gynandra* and nine other species.

Supplemental Data Set S12. Summary of repeat DNA in *G. gynandra* and *T. hassleriana*.

Supplemental Data Set S13. List of gene IDs and their RPKM values in leaf transcriptomes from S0 to S5 developmental stages of *G. gynandra* and *T. hassleriana*.

Supplemental Data Set S14. Syntenic blocks of *G. gynandra*.

Supplemental Data Set S15. Pfam annotation of the expanded genes families in *G. gynandra* genome.

Supplemental Data Set S16. The hub genes of *Dof*-GRNs of *G. gynandra* and *T. hassleriana*.

Supplemental Data Set S17. Summary of differentially expressed genes after heat stress treatment at different leaf stages in *G. gynandra* and *T. hassleriana*.

Supplemental Data Set S18. Copy numbers of genes that regulate vasculature development, C_4_ photosynthesis and heat shock response in *G. gynandra* compared to *T. hassleriana*.

Supplemental Data Set S19. Statistics analysis suggesting that *T. hassleriana* may have undergone massive gene loss compared to *G. gynandra*.

Supplemental Data Set S20. Genes that regulate vasculature development, C_4_ photosynthesis and heat shock response were mostly derived from WGD in *G. gynandra*.

Supplemental Data Set S21. Abbreviations of genes and protein enzymes in this study.

## ACKNOWLEDGEMENTS

We thank Prof. Y. Jiao at Henan Agricultural University and Prof. S. Gao at Henan University of Science and Technology for providing G. gynandra and T. hassleriana seeds used in this study. The bioinformatic analyses are performed with the supercomputing system in the Supercomputing Center of Oil Crops Research Institute, Chinese Academy of Agricultural Sciences.

## AUTHOR CONTRIBUTIONS

W.H., J.Liu, and W.Z. conceived project. W.H., J.Liu, and W.Z. designed study. J.Li, X.S. and Q.Z. conducted experiments. W.Z. and J.Liu performed data analysis. W.Z. and J.Liu wrote and revised the manuscript.

